# Bijections between the multifurcating unlabeled rooted trees and the positive integers

**DOI:** 10.1101/2023.04.19.537492

**Authors:** Alessandra Rister Portinari Maranca, Noah A. Rosenberg

## Abstract

Colijn and Plazzotta (*Systematic Biology* 67:113-126, 2018) described a bijective scheme for associating the unlabeled bifurcating rooted trees with the positive integers. In mathematical and biological applications of unlabeled rooted trees, however, nodes of rooted trees are sometimes multifurcating rather than bifurcating. Building on the bijection between the unlabeled bifurcating rooted trees and the positive integers, we describe bijective schemes for associating the unlabeled *multifurcating* rooted trees with the positive integers. We devise bijections with the positive integers for a set of trees in which each non-leaf node has *exactly k* child nodes, and for a set of trees in which each non-leaf node has *at most k* child nodes. The calculations make use of Macaulay’s binomial expansion formula. The generalization to multifurcating trees can assist with the use of unlabeled trees for applications in evolutionary biology, such as the measurement of phylogenetic patterns of genetic lineages in pathogens.

**Mathematics subject classification (2020):** 05C05, 05C30, 92D15

## 1 Introduction

In mathematical and statistical phylogenetics, the properties of evolutionary trees are used to make inferences about the processes that have given rise to those trees [2], [11]. Each tree examined in a data set is an element of some class of trees, and the mathematics pertaining to that class suggests quantities that can be informatively measured for the trees contained in the class. Often, for a label set *X* containing *n* labels, the class of trees of interest is the set of bifurcating *labeled topologies* associated with the label set. For example, if an investigator seeks to understand the evolutionary relationships among *n* named species in a taxonomic group, the phylogenetic problem of interest is to identify from data the appropriate labeled topology among the (2*n* − 3)!! possibilities.

For many phylogenetic problems, it is not the labels of specific lineages that are of interest, but rather, the *shape* of the evolutionary tree [6], [9]. For example, does the tree shape fit a model in which each lineage is equally likely to be the next to split into descendant lineages? If not, what biological features of lineages produce differences in the rates at which lineages split? Are the rates affected by abiotic factors, such as features of the environments that the lineages inhabit? For such questions, it is not the labeled topologies that are of interest, but rather, the *unlabeled* topologies. The importance of the unlabeled topologies has recently grown with the proliferation of phylodynamic studies—in which fast-evolving genetic sequences from pathogens are investigated using evolutionary trees [4], [7]. When many genetic sequences are collected across large numbers of hosts on influenza, SARS-CoV-2, or other pathogens, the placement of specific sequences in the tree is of less interest than what the tree shape reveals about transmission chains, clusters of epidemiologically related cases, and the processes that spread the pathogen. Hence, although unlabeled topologies have long been examined as mathematical objects [2], [11], phylodynamic studies have given rise to new quantities that can be computed from them as statistics for measuring their properties [1], [5].

Colijn & Plazzotta [1] introduced a variety of statistics for unlabeled topologies — bifurcating unlabeled rooted trees — focusing on metrics for comparing pairs of unlabeled topologies. Underlying some of their statistics is a bijection they devised between the set of unlabeled topologies with *n* 1 leaves and the positive integers. Each unlabeled topology *t* is associated with a positive integer that is computed recursively from the integers associated with the immediate subtrees of its root. In reverse, given an integer, the left and right subtrees associated with that integer are calculated, uniquely identifying the associated unlabeled topology. The “distance” between unlabeled topologies is the absolute difference between their associated integers. Rosenberg [8] then studied the bijection between unlabeled topologies and positive integers, characterizing the unlabeled topologies associated with the smallest and largest values among the integers associated with unlabeled topologies of *n* leaves, and determining the asymptotic growth of those quantities.

The Colijn–Plazzotta bijection and distance metric assume bifurcating trees. However, in scenarios in which the rate of divergence of evolutionary lineages is large compared to the rates of evolution along lineages individually, it is natural to instead consider multifurcating trees. In such cases, a sequence of bifurcations might be indistinguishable from a multifurcation event if insufficient time has occurred between bifurcations for mutations to accumulate. Additionally, in cases in which multiple genealogical lineages of interest trace to a single parent — such as in an instantaneous diversification of a lineage into multiple descendants — a tree has a genuine multifurcation.

Here, we introduce bijections between unlabeled *multifurcating* topologies and positive integers. We consider multifurcating trees in two different ways. First, for fixed *k* 2, we consider the class of unlabeled multifurcating topologies in which each internal node possesses exactly *k* immediate descendants (*strict k*-furcation). Next, for fixed *k* 2, we consider the class of unlabeled multifurcating topologies in which internal nodes can vary in their numbers of immediate descendants, with each node possessing at least two and *at most k* immediate descendants. The bijections between unlabeled multifurcating topologies and positive integers can be used in comparing trees in a manner analogous to use by Colijn & Plazzotta [1] of the bijection for bifurcating topologies.

In Section 2, we recall the Colijn-Plazzotta ranking scheme for bifurcating trees. Next, in Section 3, as a prelude to the general case of strictly *k*-furcating trees, we extend the scheme to strictly trifurcating trees; in Section 4, we complete the generalization. In Section 5, we consider trees that are at-most-*k*-furcating. We conclude with a discussion in Section 6.

## 2 Bifurcating trees

We begin by recalling the Colijn–Plazzotta scheme for assigning ranks to bifurcating unlabeled rooted trees [1], [8]. Let *B*_*n*_ be the set of bifurcating unlabeled rooted trees with exactly *n* leaves.

For *n* = 1, *B*_*n*_ has a single unlabeled tree with one leaf. Let 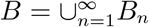 be the set of bifurcating unlabeled rooted trees, considering all possible numbers of leaves. We write *B*^∗^ = *B*\*b*_1_.

For a tree *b* ∈ *B*, let *m* : *B* → ℤ^+^ be the function that yields the number of leaves of *b*. Let *s* : *B*^∗^ → *B × B* be a function that extracts a vector containing the immediate subtrees of the root of a tree (in a canonical order that we describe shortly). We abbreviate by *b*_1_ and *b*_2_ the first and second coordinates of *s*(*b*).

### Definition 2.1.

The Colijn–Plazzotta ranking *f* : *B* → ℤ^+^ for bifurcating unlabeled rooted trees is a function that satisfies

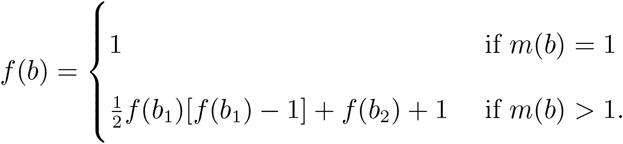

We shall abbreviate the Colijn–Plazzotta ranking as the CP ranking. To determine the CP rank of a tree, we require it to be written in a canonical form in which *f* (*b*_1_) ⩾ *f* (*b*_2_). The 1-leaf tree has CP rank 1; hence, if it is an immediate subtree of the root of *b* and *m*(*b*) ⩾ 3, then because it has the smallest CP rank among all trees, it is necessarily in the second coordinate of *s*(*b*). For convenience, we draw all trees *b* such that *b*_1_ is the left-hand subtree and *b*_2_ is the right-hand subtree. This notation will be generalized for larger trees.

That Definition 2.1 describes a bijection and yields a inverse function is proven by Colijn & Plazzotta [1] and Rosenberg [8]. For *v* = 1, the inverse function *f* ^−1^ : ℤ^+^ → *B* is the tree with one leaf, and for *v* ⩾ 2, *f* ^−1^(*v*) is the tree in *B* whose two subtrees have ranks

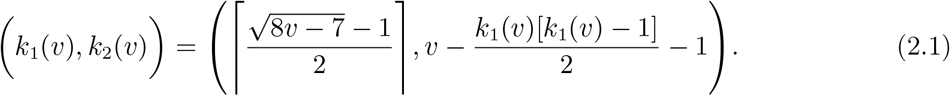

We can also examine the function *m* : *B* → ℤ^+^ that gives the number of leaves in the tree with specified CP rank. The function *m* satisfies *m*(*f* ^−1^(1)) = 1 and, for *v* ⩾ 2,

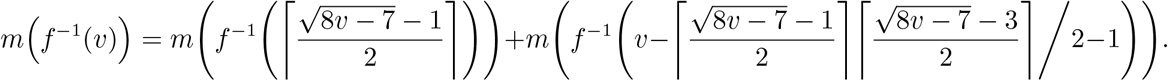

For bifurcating trees with small ranks, Figure 1 displays the trees along with their ranks and the ranks of their subtrees.

**Figure 1:**
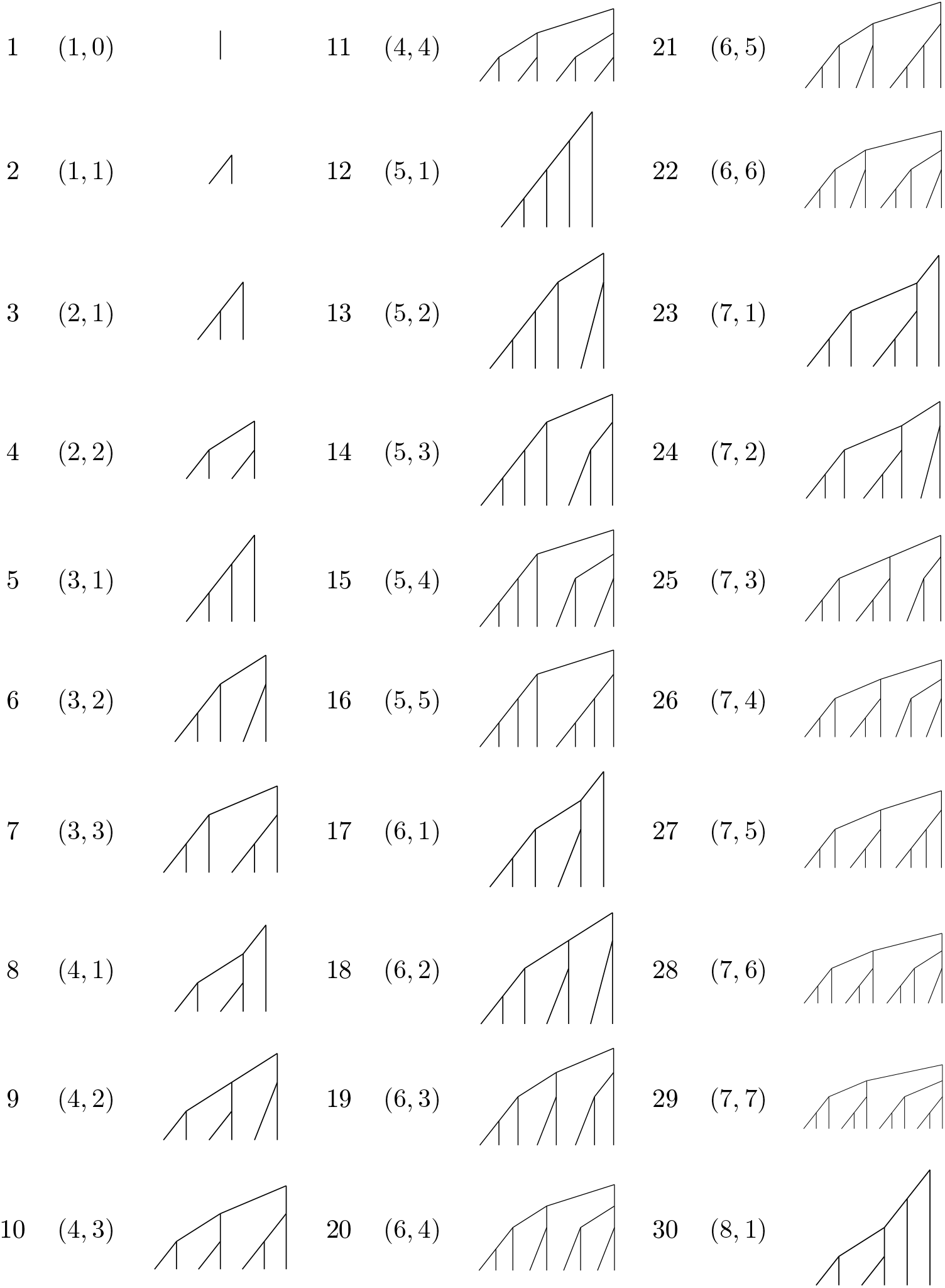
The Colijn–Plazzotta ranking for bifurcating rooted trees for small ranks. For ranks *v* from 1 to 30, the tree *b* with CP rank *v* is shown, as is the ordered pair of CP ranks of the subtrees, (*f* (*b*_1_), *f* (*b*_2_)). Each tree is drawn in canonical form, so that the rank associated with the left-hand subtree is greater than or equal to the rank associated with the right-hand subtree. The ordered pair *k*_1_(*v*), *k*_2_(*v*) is obtained from eq. 2.1, and the tree is obtained by recursive application of eq. 2.1.

## 3 Trifurcating trees

To prepare for the general *k*-furcating case, we now consider trifurcating trees, in which each internal node possesses exactly three immediate descendant nodes. Let *T*_*n*_ be the set of trifurcating unlabeled rooted trees with exactly *n* leaves. As was true for the bifurcating case, for the trivial tree with *n* = 1, *T*_*n*_ consists of a single unlabeled tree with one leaf. Let 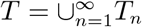 be the set of trifurcating unlabeled rooted trees, considering all possible numbers of leaves, and let *T* ^∗^ = *T \ T*1. Note that for even values of *n, T*_*n*_ is empty, as no strictly trifurcating tree can possess an even number of leaves. We continue to use the notation *m* and *s* for concepts analogous to those of the bifurcating case.

For a tree *t* ∈ *T*, let *m* : *T* → ℤ^+^ count the number of leaves of *t*. Let *s* : *T* ^∗^ → *T* × *T* × *T* denote the vector that extracts the immediate subtrees of the root of a tree. We abbreviate by *t*_1_, *t*_2_, *t*_3_ the first, second, and third coordinates of *s*(*t*), in a canonical order discussed below.

### 3.1 Ranking scheme

We seek to find a bijection *f* : *T* → ℤ^+^ between trifurcating trees and positive integers. We accomplish this task in a manner analogous to that used in the bifurcating case. In that case, we devised a polynomial that is quadratic in the rank of the first subtree and linear in that of the second subtree. With three subtrees, we increase the degree of the polynomial by one, and compute a cubic term for the first subtree, a quadratic term for the second, and a linear term for the third. Consider a tree *t* = (*t*_1_, *t*_2_, *t*_3_). We assume a canonical form in which *f* (*t*_1_) ⩾*f* (*t*_2_) ⩾ *f* (*t*_3_); this form places a dictionary order on trees. To assign the rank *f* (*t*), we must sum three quantities: (1) the number of nontrivial trees in *T* whose first subtree has rank less than *f* (*t*_1_); (2) the number of nontrivial trees in *T* whose first subtree has rank *f* (*t*_1_) and second subtree has rank less than *f* (*t*_2_); and (3) the number of nontrivial trees in *T* whose first subtree has rank *f* (*t*_1_), second subtree has rank *f* (*t*_2_), and third subtree has rank less than *f* (*t*_3_). We then assign *t* the rank equal to the sum of the quantities in (1), (2), and (3), plus 2. The +2 assigns the next available rank to *t*, accounting for the trivial tree with *n* = 1; we can view the trivial tree as having subtrees with ranks (1, 0, 0).

For quantity (3), the number of nontrivial trees in *T* whose first subtree has rank *f* (*t*_1_), whose second tree has rank *f* (*t*_2_), and whose third subtree has rank less than *f* (*t*_3_), we count all trees with subtrees (*t*_1_, *t*_2_, *y*_3_), where *y*_3_ ranges from 1 to *f* (*t*_3_) − 1. The number of such trees is

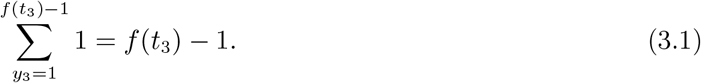

For quantity (2), we count nontrivial trees in *T* whose first subtree has rank *f* (*t*_1_) and whose second tree has rank less than *f* (*t*_2_). The number of such trees is

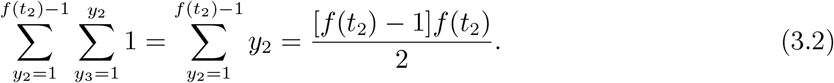

Finally, for quantity (1), we count nontrivial trees in *T* with first subtree rank less than *f t*1):

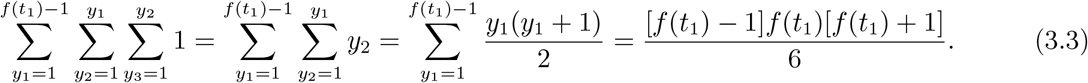

We sum eqs. 3.3, 3.2, and 3.1, plus 2, and we obtain the index for tree *t*:

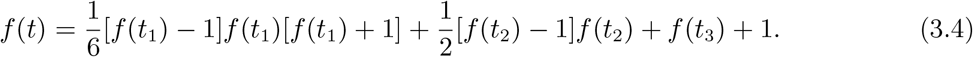

As an example, consider tree *t* = (*t*_1_, *t*_2_, *t*_3_) with (*f* (*t*_1_), *f* (*t*_2_), *f* (*t*_3_)) = (4, 3, 3). With eq. 3.4, we Obtain 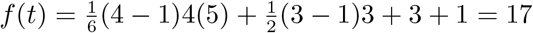. Hence, the tree with subtree ranks (4, 3, 3) is the tree with rank 17.

#### Definition 3.1.

The ranking *f* : *T* → ℤ^+^ for trifurcating unlabeled rooted trees is a function that satisfies

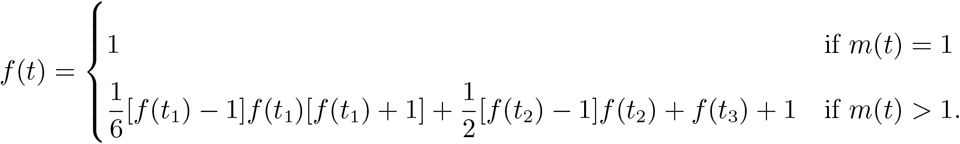

For trifurcating trees with small ranks, Figure 2 displays the trees along with their ranks and the ranks of their subtrees.

**Figure 2:**
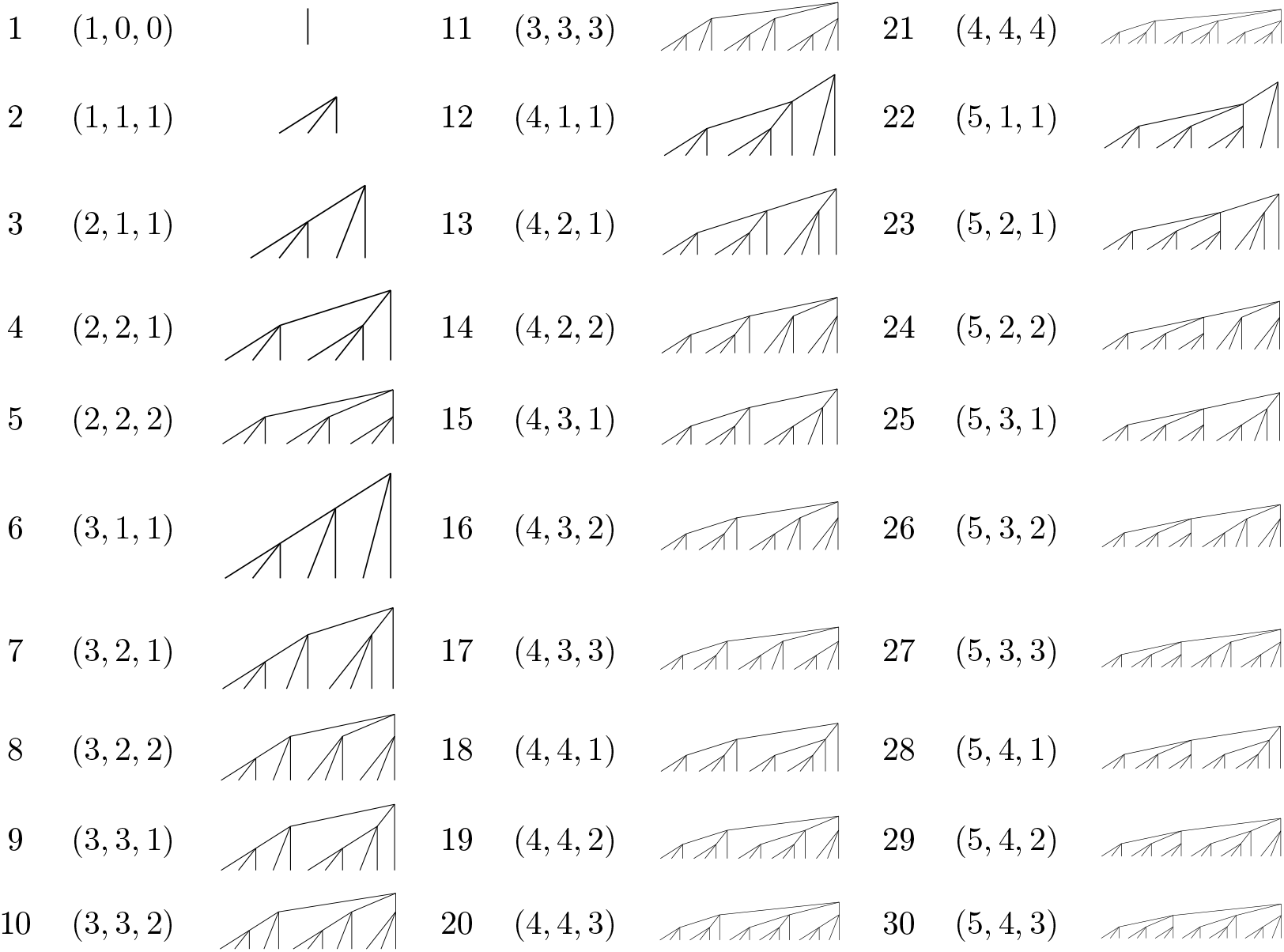
Trifurcating trees associated with specified ranks. For each rank *v* from 1 to 30, the ranks The ordered triple (*k*_1_(*v*), *k*_2_(*v*), *k*_3_(*v*)) is obtained from Theorem 3.3, and the tree is obtained by recursive *k*_1_(*v*), *k*_2_(*v*), *k*_3_(*v*) (of its three subtrees appear, followed by the trifurcating tree associated with rank *v*. application of Theorem 3.3.

### 3.2 Bijectiveness of ranking scheme

That our ranking for trifurcating trees bijectively associates *T* with the natural numbers ℤ^+^ is clear by construction. As in the Colijn-Plazzotta ranking for bifurcating trees, underlying the construction is the idea that given a tree *t*, the integer assigned to *t* is obtained by counting trees whose ranks are less than those of *t* and adding 1 to yield the rank of *t*. Trees are ordered lexicographically, as illustrated in Figure 2. Each positive integer is trivially assigned a rank (surjectivity). That two distinct trees *t* and *t*^*i*^ possess different ranks is likewise trivial, as one of the two trees must be enumerated among the trees with lower ranks than the other.

Nevertheless, an algebraic proof that our ranking for trifurcating trees bijectively associates *T* with the natural numbers ℤ^+^ is instructive. The proof illustrates the way in which the three subtree ranks are analogous to the elementary concept of “place value.” Each subtree entry is associated with a polynomial: cubic for the first, quadratic for the second, and linear for the third. As in “place value,” the contribution to the rank from each of the latter two entries is bounded above by the increase in rank caused by the smallest possible increment to its predecessor.

#### Theorem 3.2.

The function *f* : *T* → ℤ^+^ is a bijection.

*Proof*. To prove injectivity, first note that for the trivial tree, (*f* (1, 0, 0)) = 1, and for any tree *t* ≠ (1, 0, 0), *f* (*t*) > 1. Next, we must show that for two distinct nontrivial trees *t* = (*t*_1_, *t*_2_, *t*_3_) and *y* = (*y*_1_, *y*_2_, *y*_3_), *f* (*t*) ≠ *f* (*y*). We can separate the problem into three cases: (i) *t*_1_ = *y*_1_, *t*_2_ = *y*_2_, and *t*_3_ ≠ *y*_3_; (ii) *t*_1_ = *y*_1_ and *t*_2_ ≠ *y*_2_; (iii) *t*_1_ ≠ *y*_1_.

Case (i). Suppose *t*_1_ = *y*_1_ and *t*_2_ = *y*_2_; without loss of generality, assume *f* (*t*_3_) > without loss of generality, assume *f* (*y*_3_). Then *f* (*t*) − *f* (*y*) = *f* (*t*_3_) − *f* (*y*_3_) > 0,

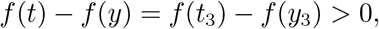

so that *f* (*t*) ≠ *f* (*y*).

Case (ii). Suppose *t*_1_ = *y*_1_, and without loss of generality, assume *f* (*t*_2_) > *f* (*y*_2_). Then

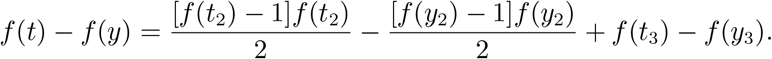

Because *f* (*t*_2_) > *f* (*y*_2_) and both *f* (*t*_2_) and *f* (*y*_2_) are integers, *f* (*t*_2_) ⩾ *f* (*y*_2_) + 1, and

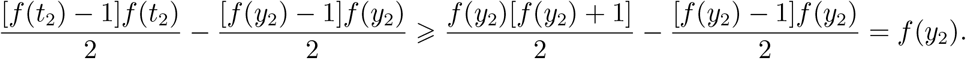

Next, note that because *t* is not the trivial tree (1, 0, 0), *f* (*t*_3_) ⩾ 1. Because a tree is required to be in canonical form, *f* (*y*_3_) ⩽ *f* (*y*_2_). Hence *f* (*t*_3_) − *f* (*y*_3_) ⩾ 1 − *f* (*y*_2_). Then *f* (*t*) − *f* (*y*) ⩾ *f* (*y*_2_) +(1 − *f* (*y*_2_)) = 1, and *f* (*t*) ≠ *f* (*y*).

Case (iii). Suppose *t*_1_ ≠ *y*_1_. We can assume without loss of generality that *f* (*t*_1_) > *f* (*y*_1_). Then

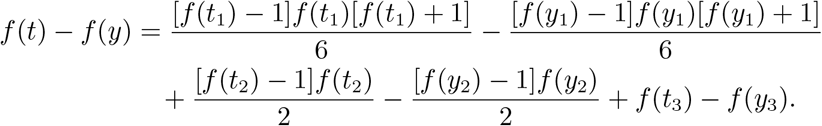

Because *f* (*t*_1_) > *f* (*y*_1_), and both *f* (*t*_1_) and *f* (*y*_1_) are integers, *f* (*t*_1_) ⩾*f* (*y*_1_) + 1, and

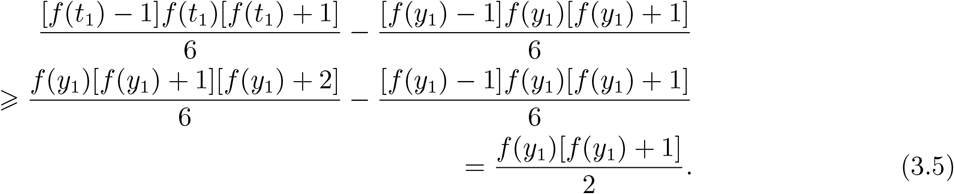

We also know that because *f* (*y*_2_) ⩽ *f* (*y*_1_),

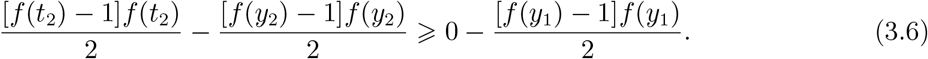

As in Case (ii), we have

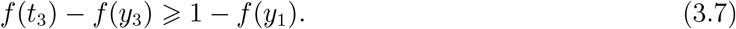

Combining inequalities 3.5, 3.6, and 3.7, we see that

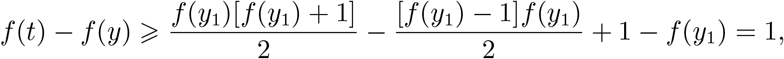

and *f* (*t*) ≠ *f* (*y*).

With all three cases demonstrated, we conclude that if (*t*_1_, *t*_2_, *t*_3_) ≠ (*y*_1_, *y*_2_, *y*_3_), then *f* (*t*) ≠ *f* (*y*), so that *f* is injective.

For surjectivity, each positive integer *v* ⩾ 2 has a unique representation as a decomposition

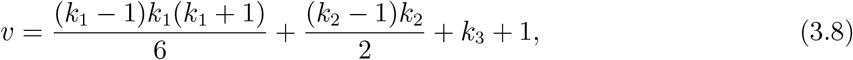

with *k*_1_, *k*_2_, *k*_3_ positive integers and *k*_1_ ⩾ *k*_2_ ⩾ *k*_3_. To understand why such a decomposition exists, note that as *k*_3_ ranges from 1 to *k*_2_, (*k*_2_ − 1)*k*_2_/2 + *k*_3_ ranges from (*k*_2_ − 1)*k*_2_/2 + 1 to *k*_2_(*k*_2_ + 1)/2, so that the ordered pairs (*k*_2_, *k*_3_) with *k*_2_ and *k*_3_ variable, *k*_1_ ⩾ *k*_2_ ⩾ *k*_3_ ⩾ 1, enumerate all positive integers from 1 to *k*_1_(*k*_1_ + 1)/2.

Next, as *k*_2_ ranges from 1 to *k*_1_ and *k*_3_ ranges from 1 to *k*_2_, (*k*_1_ − 1)*k*_1_(*k*_1_ + 1)/6 +(*k*_2_ − 1)*k*_2_/2 + *k*_3_ ranges from (*k*_1_ − 1)*k*_1_(*k*_1_ + 1)/6 + 1 to *k*_1_(*k*_1_ + 1)(*k*_1_ + 2)/6. Hence, the ordered pairs (*k*_1_, *k*_2_, *k*_3_) with *c* ⩾ *k*_1_ ⩾ *k*_2_ ⩾ *k*_3_ enumerate all positive integers from 1 to *c*(*c* + 1)(*c* + 2)/6.

Noting that 1 is added in the decomposition in eq. 3.8, eq. 3.8 traverses all the positive integers greater than or equal to 2.

### 3.3 The inverse function that recursively converts an integer to a tree

#### Theorem 3.3.

The function *f* ^−1^ : ℤ^+^ → *T* gives the three coordinates of the tree whose rank is *v*, and it satisfies

a. *f* ^−1^(1) is the tree with one leaf, and
b. for *v* ⩾ 2, *f* ^−1^(*v*) is the tree *t* ∈ *T* whose subtrees have the ranks:

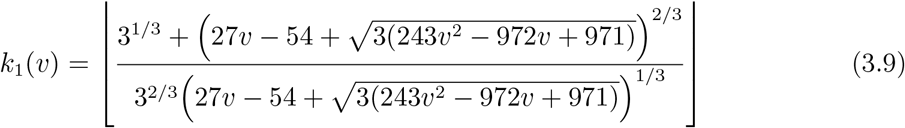

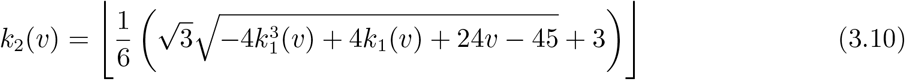

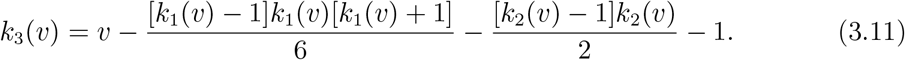

*Proof*. For *v* ⩾ 2, we find the unique (*f* (*t*_1_), *f* (*t*_2_), *f* (*t*_3_)) with *f* (*t*_1_) ⩾ *f* (*t*_2_) ⩾ *f* (*t*_3_) ⩾ 1 that solves

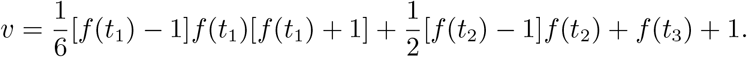

First, *f* (*t*_1_) is the largest integer satisfying

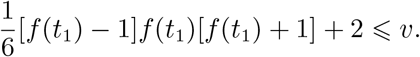

Solving the inequality, the first subtree has rank as in eq. 3.9.

Next, *f* (*t*_2_) is the largest integer satisfying

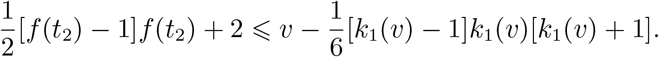

Solving, the second subtree has rank as in eq. 3.10. Finally, *t*_3_ has rank as in eq. 3.11.

Using the inverse function *f* ^−1^, Figure 2 gives the trifurcating trees for ranks 1 to 30.

### 3.4 Number of leaves associated with the tree of a given integer

With the bijection between trifurcating trees and positive integers in Theorem 3.3, we quickly obtain a recursion for the number of leaves possessed by a trifurcating tree with specified rank.

#### Theorem 3.4.

The function *m* : *B* → ℤ^+^ that gives the number of leaves in the tree *f* ^−1^(*v*) with specified rank satisfies *m* (*f* ^−1^(1)) = 1, and for *v* ⩾ 2,

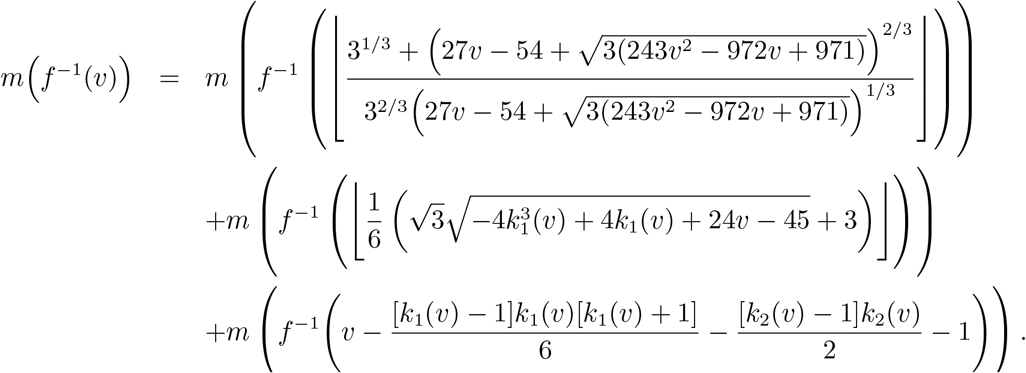

*Proof*. The number of leaves in the tree of rank *v* ⩾ 2, or *m* (*f* ^−1^(*v*)), is the sum of the numbers of leaves in its first, second, and third subtrees, or *m* (*k*_1_(*v*)) + *m* (*k*_2_(*v*)) + *m* (*k*_3_(*v*)).

## 4 *k*-furcating trees

The results for bifurcating and trifurcating trees generalize. Let *U*_*n*_ be the set of *k-furcating* unlabeled rooted trees with *n* leaves. For *n* = 1, *U*_*n*_ has a single unlabeled tree with one leaf. Suppose each non-leaf node of a tree with *n* ⩾ *k* leaves has exactly *k* descendants. Let 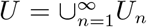 be the set of *k*-furcating unlabeled rooted trees, considering all possible numbers of leaves. Let *U* ^∗^ = *U \U_n_*. We describe a bijection between unlabeled *k*-furcating rooted trees and positive integers.

For a tree *u* ∈ *U, m* : *U* → ℤ^+^ yields the number of leaves of *u*, and *s* : *U* ^∗^ → *U* × *U* × …· × *U* extracts a vector containing the immediate subtrees of the root of a tree. We abbreviate by *U*_*n*_, *u*_2_, …, *u*_*k*_ the *k* coordinates of *s*(*u*). The case of *k* = 2 is the Colijn-Plazzotta ranking. The case of *k* = 3 is the case described in Section 3. We now consider arbitrary *k* ⩾ 2.

### 4.1 Ranking scheme

We seek to derive the rank of a tree *u* = (*U*_*n*_, *u*_2_, …, *u*_*k*_). We must sum *k* quantities: (1) the number of nontrivial trees in *U* whose first subtree has rank less than *f* (*U*_*n*_); (2) the number of nontrivial trees in *U* whose first subtree has rank *f* (*U*_*n*_) and whose second subtree has rank less than *f* (*u*_2_); … (*k*) the number of nontrivial trees whose first subtree has rank *f* (*U*_*1*_), whose second subtree has rank *f* (*u*_2_), …, whose (*k* − 1)-th subtree has rank *f* (*u*_*k*_−1), and whose *k*th subtree has rank less than *f* (*u*_*k*_). We assign to *u* the next available rank, equal to the sum of the quantities in (1), (2), …, (*k*) plus 2, accounting for the trivial tree with *n* = 1.

The number of trees whose first term is less than *f* (*U*_*1*_) is the number of trees satisfying *f* (*U*_*1*_) > *f* (*u*_2_) ⩾ *f* (*u*_3_) ⩾ … ⩾ *f* (*u*_*k*_). Coordinate *f* (*u*_2_) is strictly less than *f* (*U*_*n*_), *f* (*u*_3_) can be at most *f* (*u*_2_), and so on. Because all coordinates take integer values, the desired number of trees is

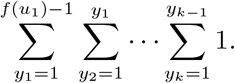

More generally, considering only the “end” of a vector of subtrees, from coordinate *n* ⩽ *k* to coordinate *k*, the number of trees whose *n*th term is less than *f* (*U*_*n*_) follows the same logic. We must consider all possible values (*u,_1_, u_n_*+1, …, *u*_*k*_) with *f* (*U*_*n*_+1) ⩽ *f* (*U*_*n*_) − 1, *f* (*U*_*n*_+2) ⩽ *f* (*U*_*n*_+1), and so on, so that the desired number of trees is

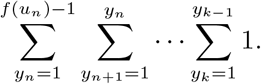

From this expression, we devise a formula for the rank of a tree *u*.

We have the following lemma.

#### Lemma 4.1.

The number of nonincreasing sequences of positive integers (*y*_*n*_, *y*_*n+1*_, …, *y*_*k*_) in which all entries are bounded above by *y*_*n*_−1 is

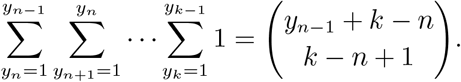

*Proof*. We proceed by induction, working from inner sums outward. For the inner sum, with *n*

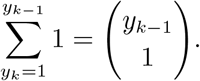

Now assume that our statement holds for the sum indexed by *y*_*n+1*_. That is, assume that

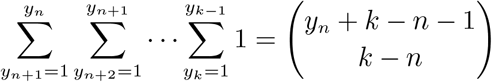

Therefore,

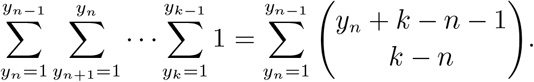

Using the standard combinatorial identity for positive integers *m, n*,

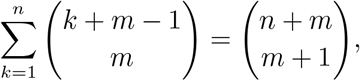

we see that, as desired,

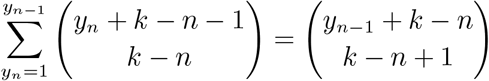

#### Definition 4.2.

Consider a *k*-furcating tree *u* = (*U*_*n*_, *u*_2_, …, *u*_*k*_). The ranking *f* : *U* → ℤ^+^ for *k*-furcating unlabeled rooted trees is a function that satisfies *f* (*u*) = 1 for the one-leaf tree with *m*(*u*) = 1, and for *m*(*u*) > 1,

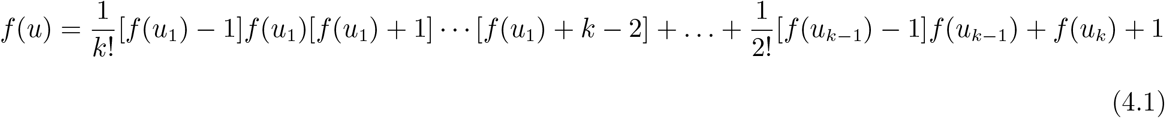

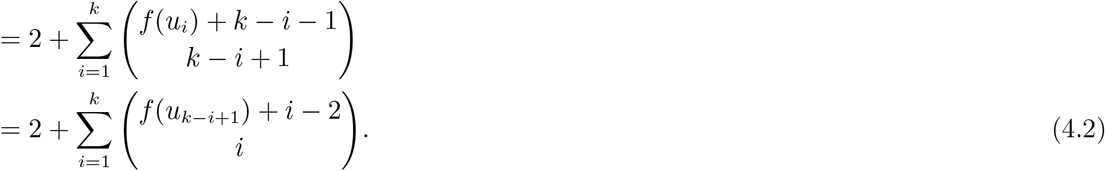

To obtain this definition, we have applied Lemma 4.1 term-wise to each of the *k* entries of the vector of subtrees associated with tree *u*. If *f* (*u*_*k*_) + 1 is more instructively written [*f* (*u*_*k*_) − 1] + 2, then the term with index *i* in eq. 4.1 counts the number of trees with fixed rank in preceding positions and rank less than *f* (*u*_*i*_) in position *i*; we adjust the sum by +2 to assign the next available rank.

For *k* = 3, Definition 4.2 reduces to Definition 3.1; for *k* = 2, it reduces to Definition 2.1.

### 4.2 Bijectiveness of ranking scheme

As in the bifurcating and trifurcating cases, that Definition 4.2 bijectively relates multifurcating rooted trees and positive integers is clear from the construction. An algebraic argument demonstrating the bijection can make use of Macaulay’s binomial expansion theorem [10], [12].

#### Theorem 4.3.

(Macaulay’s binomial expansion theorem [10]) Each positive integer can be decomposed as a certain sum of binomial coefficients. In particular, let *h* and *i* be positive integers. Then *h* can be uniquely written in the form

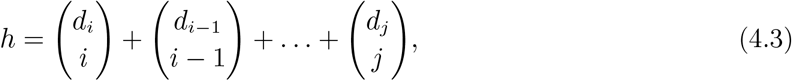

where *d*_*k*_ takes integer values for all *k* and *d*_*i*_ > *d*_*i*_−1 > … > *dj* ⩾ *j* ⩾ 1. The expression in eq. 4.3 is called the *i*-binomial expansion of the integer *h*.

We can make a slight adaptation of Macaulay’s binomial expansion theorem. By the theorem, for fixed *i* > 0, each integer *h* > 0 can be uniquely written in the form

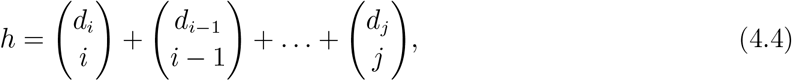

where *d*_*k*_ takes integer values for all *k* and *d*_*i*_ > *d*_*i*_−1 > … > *dj*−1 ⩾ *j* ⩾ 1, and *dj* ⩾ 0. We restate this adaptation as a corollary to Theorem 4.3.

#### Corollary 4.4.

If the last entry in the decreasing sequence (*d*_*i*_, *d*_*i*_−1, …, *dj*) is permitted to equal 0, then the decomposition of *h* in the form of eq. 4.4 is still unique.

*Proof*. We take the *i*-binomial expansion of the integer *h* according to Theorem 4.3. If it does not contain a term 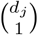, then we append a term 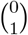 to make a modified *i*-binomial expansion.

#### Theorem 4.5.

The function *f* : *U* → ℤ^+^ described in Definition 4.2 is bijective.

*Proof*. Definition 4.2 associates the 1-leaf tree with *f* (*u*) = 1 and the *k*-furcating tree of *k* leaves with *f* (*u*) = 2. We wish to show that for all other *k*-furcating trees *u*, the value produced by Definition 4.2 for tree *u* is greater than or equal to 3 and is associated only with tree *u* (injectivity). We also wish to show that each positive integer greater than or equal to 3 is associated with a tree *u* (surjectivity).

For injectivity, in our Definition 4.2, for each *i* from 1 to *k*, let

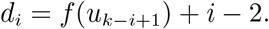

Then eq. 4.2 can be rewritten

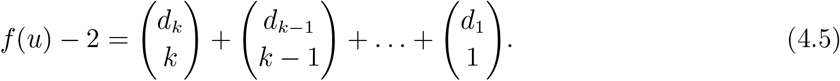

We have *d*_*i*_+1 > *d*_*i*_; we see this by noting that *f* (*u*_*k*_−(*i*+1)+1) ⩾ *f* (*u*_*k*_−*i*+1), so that

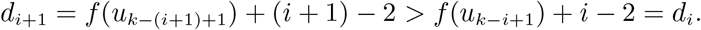

We can now make use of the modification of Macaulay’s binomial expansion theorem in Corollary 4.4, applied with the integer *f* (*u*) − 2 in the role of *h* and *k* in the role of *i*. The modified theorem states that *f* (*u*) − 2 has a unique modified *k*-binomial decomposition; as eq. 4.5 describes such a decomposition, that decomposition is unique.

For surjectivity, each positive integer *f* (*u*) ⩾ 3 can be applied in eq. 4.5 to recover the associated (*f* (*U*_*n*_), *f* (*u*_2_), …, *f* (*u*_*k*_)).

### 4.3 The inverse function that recursively converts an integer to a tree

We can apply the bijective representation of *k*-furcating trees with positive integers to identify the tree assocciated with a specific integer. With the exception of the 1-leaf tree, associated with the integer 1, a *k*-furcating tree has the form (*U*_*n*_, *u*_2_, …, *u*_*k*_).

The function *f* ^−1^ : ℤ^+^ → *U* gives a tree; to obtain *f* ^−1^(*v*) for a tree with rank *v*, we first compute the vector of ranks of the immediate subtrees of the root of the tree with rank *v*:

a. *f* ^−1^(1) is the tree with one leaf.
b. For *v* ⩾ 2, *f* ^−1^(*v*) is the tree *u* ∈ *U* whose subtrees have the ranks given by the algorithm below.

1. To find the first term, *f* (*U*_*n*_) is the largest integer satisfying

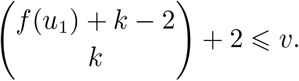
2. As in the 3-furcating case, we remove the contributions to the total rank of previous elements. Proceeding sequentially from *j* = 2 until *j* = *k*, the *j*th term in (*f* (*U*_*n*_), *f* (*u*_2_), …, *f* (*u*_*k*_)) is obtained from terms 1 to *j* − 1 by computing the largest integer satisfying

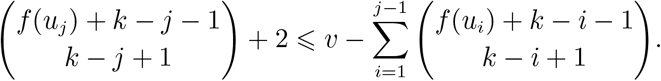

The algorithm gives the ranks of the *k* immediate subtrees of the root of the tree with rank *v*; to obtain the *entire* tree for rank *v*, we apply the algorithm recursively to ranks *f* (*U*_*n*_), *f* (*u*_2_), …, *f* (*u*_*k*_). Note that in order to calculate the number of leaves associated with the tree of rank *v*, we recursively proceed by identifying the *k* subtrees of tree *f* ^−1^(*v*), summing their numbers of leaves.

In other words, the function *m* : *U* → ℤ^+^ that gives the number of leaves in the tree with specified rank *v* satisfies *m* (*f* ^−1^(1)) = 1, and for *v* ⩾ 2,

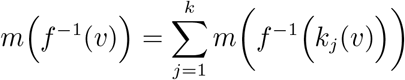

with *kj*(*v*) representing the rank of the *j*th coordinate in the tree with rank *v*, as obtained when the algorithm proceeds through index *j*.

As an example, Figure 3 gives the bijection for 4-furcating trees associated with ranks 1 to 30.

**Figure 3:**
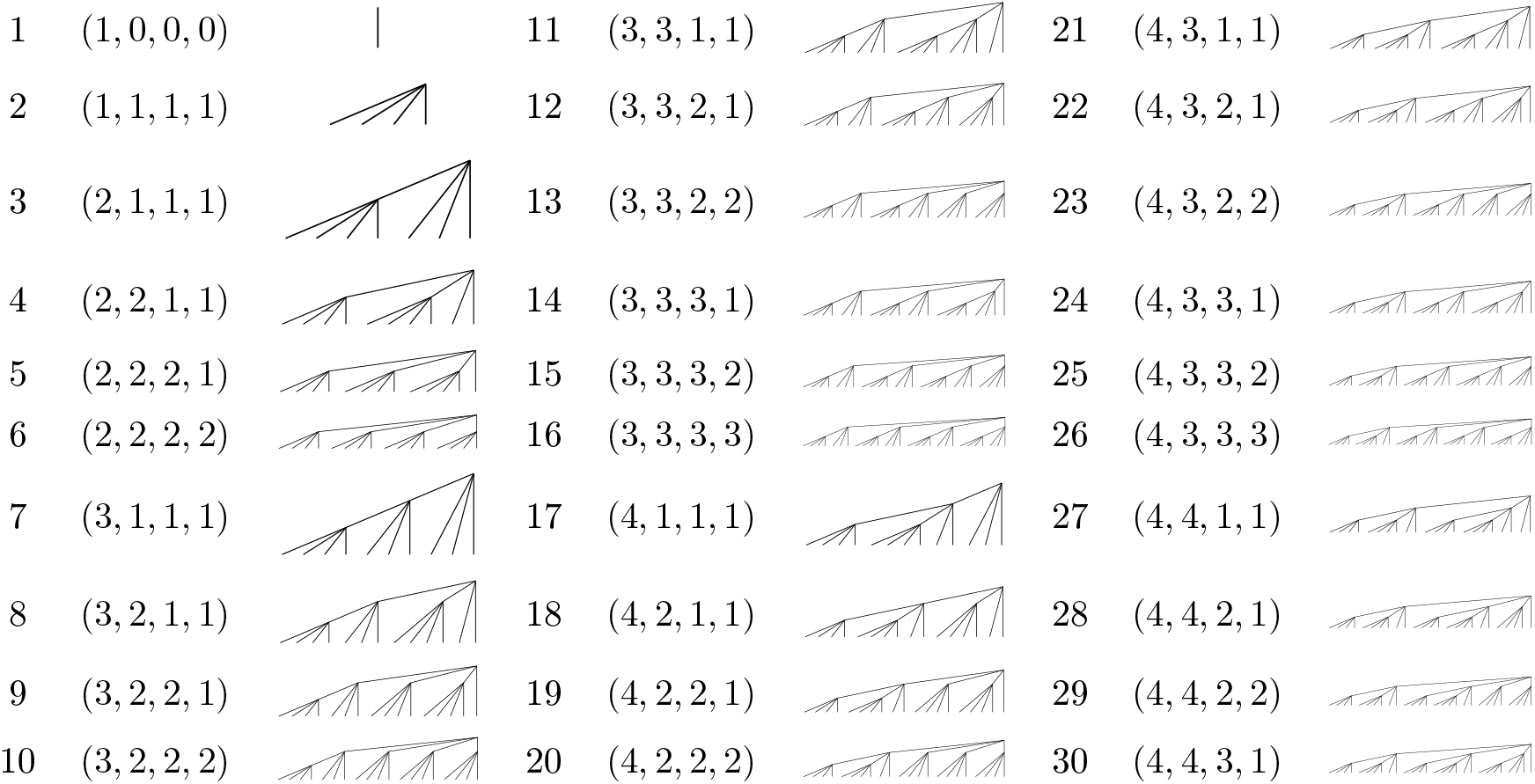
4-furcating trees associated with specified ranks. For each rank *v* from 1 to 30, the ranks The ordered quadruple (*k*_1_(*v*), *k*_2_(*v*), *k*_3_(*v*), *k*4(*v*)) is obtained from the algorithm in Section 4.3, and the tree *k*_1_(*v*), *k*_2_(*v*), *k*_3_(*v*), *k*4(*v*() of its four subtrees appear, followed by the 4-furcating tree associated with rank *v*. is obtained by recursive application of the algorithm.

## 5 At-most-*k*-furcating trees

We next consider *at-most-k-furcating* trees, trees for which each non-leaf node possesses *at most k* descendants. Let *An* be the set of unlabeled rooted trees with *n* leaves, such that each non-leaf node possesses at most *k* descendants. Let 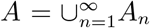 be the set of all at-most-*k*-furcating unlabeled rooted trees. We include the single-leaf tree for *n* = 1, and write *A*^∗^ = *A \ A*1.

For a tree *a* ∈ *A*, let *m* : *A* → ℤ^+^ denote the number of leaves of *a*. The function *s* : *A*^∗^ → *A* × *A* × (*A* ∪ ∅) × … × (*A* ∪ ∅) contains the immediate subtrees of the root of a tree. Because trees in *A* are *at-most k*-furcating, each non-leaf node possesses at least two and at most *k* descendants; in a canonical ordering of the subtrees of a non-leaf node, subtrees 3, 4, …, *k* can be empty.

We now describe a bijection between unlabeled at-most-*k*-furcating rooted trees and positive integers. Note that as before, the case of *k* = 2 is the Colijn-Plazzotta ranking; *k* = 3 is the smallest case for which the sets of strictly *k*-furcating trees and at-most-*k*-furcating trees differ.

### 5.1 Ranking scheme

Our goal is to obtain a rank *f* (*a*) for a tree *a* = (*a*_1_, *a*_2_, …, *a*_*k*_). We make use of a standard combinatorial identity. In particular, for integers *m, n* ⩾ 0

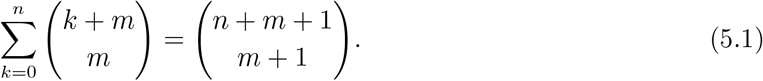

We will also use the following lemma.

#### Lemma 5.1.

The number of non-increasing sequences of nonnegative integers (*yn, yn*+1, …, *y*_*k*_) in which all entries are bounded above by *y*_*n*_−1 is

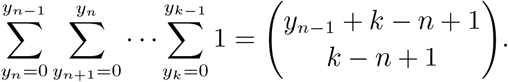

*Proof*. We proceed by induction, working from the innermost sum outward. First, consider the innermost sum, with *n* = *k*:

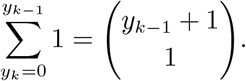

Now assume that our statement holds for the sum indexed by *y*_*n*_+1. That is, assume that

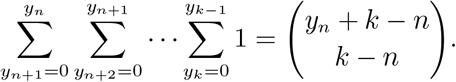

Therefore,

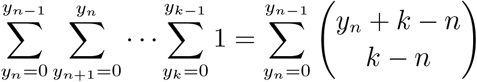

Using the combinatorial identity in eq. 5.1,

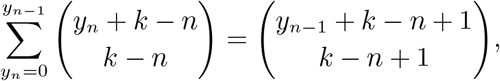

as desired.

As in Section 4.2, to find the rank of *a*, we must sum (1) the number of nontrivial trees in *A* whose first subtree has rank less than *f* (*a*_1_); (2) the number of nontrivial trees in *A* whose first subtree has rank *f* (*a*_1_) and whose second subtree has rank less than *f* (*a*_2_); (3) the number of nontrivial trees in *A* whose first subtree has rank *f* (*a*_1_), second subtree has rank *f* (*a*_2_), and third subtree has rank less than *f* (*a*_3_). Here, unlike in the strictly *k*-furcating case, we allow the third subtree to be empty. Continuing, with subsequent subtrees, the last item in the sum is the number of nontrivial trees in *A* whose first subtree has rank *f* (*a*_1_), second subtree has rank *f* (*a*_2_), …, (possibly empty) (*k* − 1)th subtree has rank *f* (*a*_*k*_−1) and (possibly empty) *k*th subtree has rank less than *f* (*a*_*k*_). We assign *a* the next available rank, which, accounting for the trivial tree with *n* = 1, is 2 more than the quantities we have summed. The required sum is

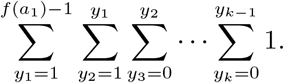

Note that the two outermost sums begin from 1, as these terms correspond to nonempty subtrees; the remaining *k* − 2 indices are permitted to equal 0.

Using eq. 5.1 and Lemma 5.1, the number of trees with second subtree rank less than a fixed second term *f* (*a*_2_) is given by

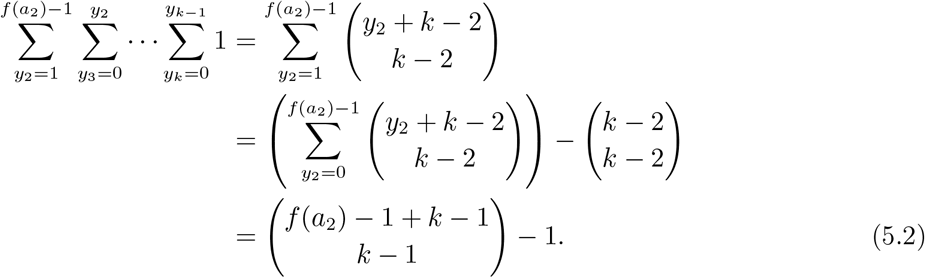

Next, to find the number of trees with rank less than the first term *f* (*a*_1_), we have

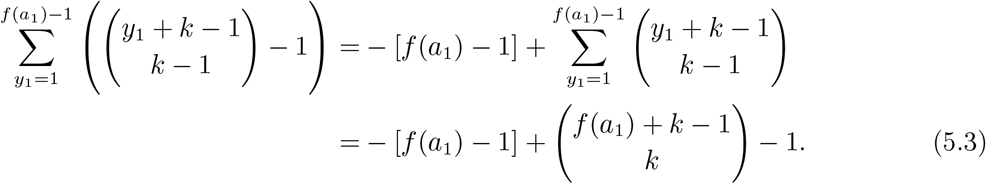

Finally, for a general fixed term *f* (*an*), we have, analogously to the *k*-furcating case, the number of trees with *n*th subtree rank less than the fixed term *f* (*an*) is given by

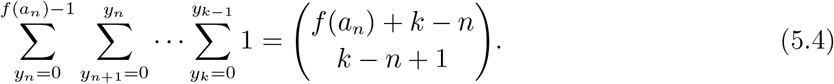

Eqs. 5.2, 5.3 and 5.4 yield our desired ranking, as described in the following definition. **Definition 5.2**. Consider an at-most-*k*-furcating tree *a* = (*a*_1_, *a*_2_, …, *a*_*k*_). The ranking *f* : *A* → ℤ^+^ for at-most-*k*-furcating unlabeled rooted trees is a function that satisfies *f* (*a*) = 1 for the one-leaf tree with *m*(*a*) = 1, and, for *m*(*a*) > 1,

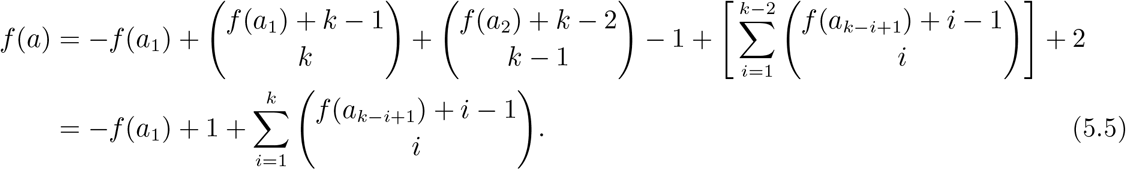

Here, we have applied eq. 5.1 to each of the *k* entries of the vector of subtrees for tree *a*. For *k* = 2, Definition 5.2 reduces to Definition 2.1.

### 5.2 Bijectiveness of ranking scheme

As in the case of strictly *k*-furcating trees, the construction that gives rise to Definition 5.2 provides a bijective ranking scheme between at-most-*k*-furcating trees and positive integers. We state the result for completeness but omit a detailed algebraic argument.

#### Theorem 5.3.

The function *f* : *A* → ℤ^+^ as defined in Definition 5.2 is a bijection.

### 5.3 The inverse function that recursively converts an integer to a tree

As in the strictly *k*-furcating case, we can obtain the at-most-*k*-furcating tree associated with a specified integer. An at-most-*k*-furcating tree has the form (*a*_1_, *a*_2_, …, *a*_*k*_), with terms after *a*_2_ possibly empty. The function *f* ^−1^ : ℤ^+^ → *T* gives the tree whose rank is *v*:

a. *f* ^−1^(1) is the tree with one leaf, and
b. for *v* ⩾ 2, *f* ^−1^(*v*) is the tree *t* ∈ *T* whose subtrees have the ranks given by the algorithm below.

1. To find the first term, *f* (*a*_1_) is the largest integer satisfying

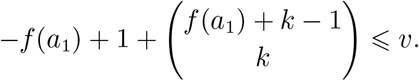
2. Proceeding sequentially from *j* = 2 until *j* = *k*, the *j*th term in (*f* (*a*_1_), *f* (*a*_2_), …, *f* (*a*_*k*_)) is obtained from terms 1 to *j* − 1 by computing the largest integer satisfying

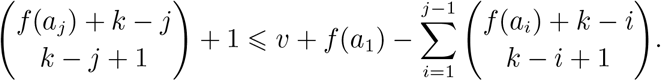

After step 2, all *k* coordinates are obtained for the rank *v*. Note that step 2 can recover values of 0 for coordinates after the first two. We then have that the function *m* : *A* → ℤ^+^ that gives the number of leaves in the tree with specified rank satisfies *m* (*f* ^−1^(1)) = 1, and for *v* ⩾ 2,

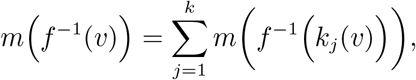

with *kj*(*v*) representing the rank of the *j*th coordinate in the tree with rank *v*, as obtained when the algorithm proceeds through index *j*.

Figure 4 provides the bijection for at-most-trifurcating trees associated with ranks 1 to 30, and Figure 3 gives the corresponding bijection for at-most-4-furcating trees.

**Figure 4:**
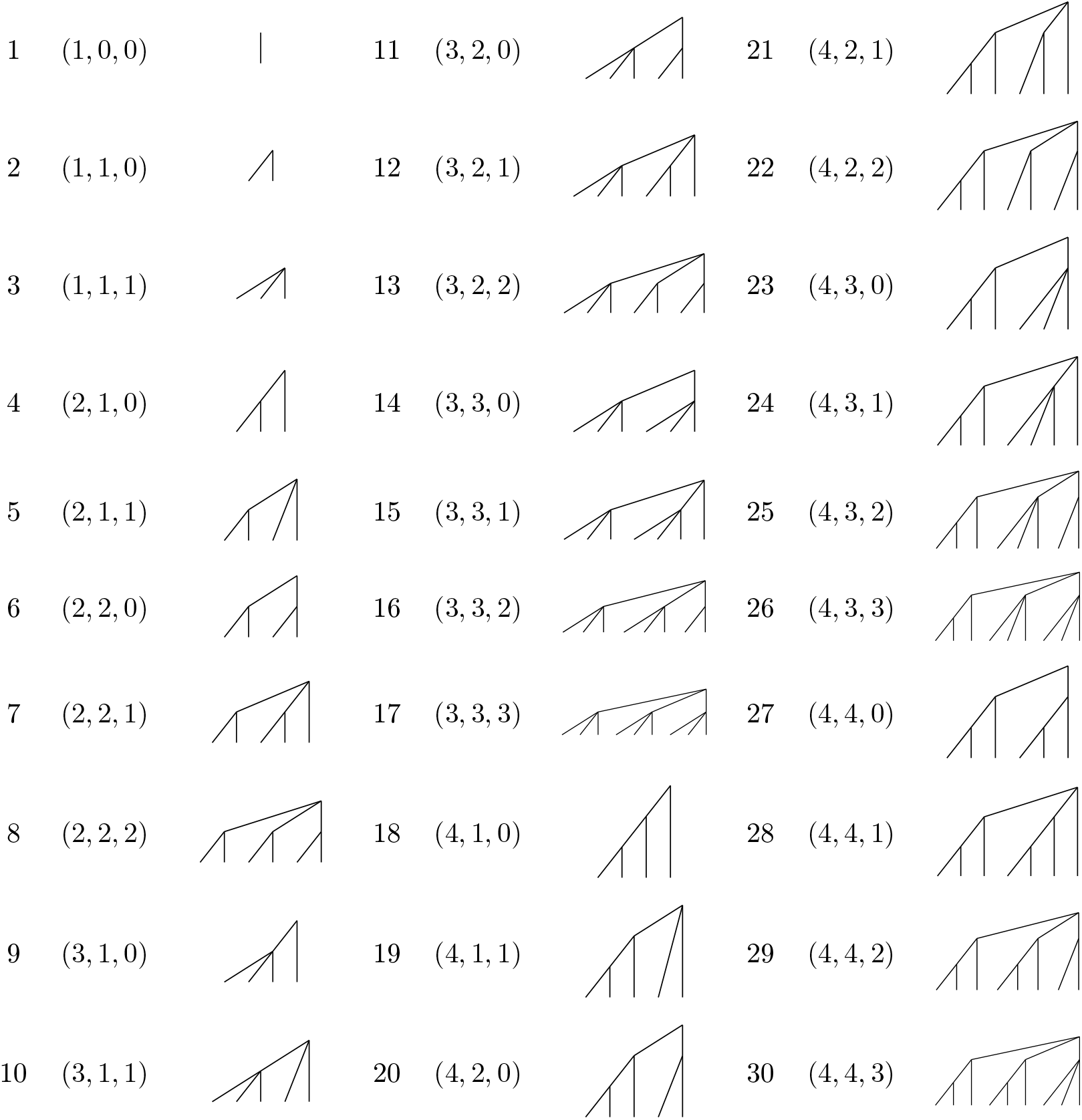
At-most-trifurcating trees associated with specified ranks. For each rank *v* from 1 to 30, the ranks The ordered triple (*k*_1_(*v*), *k*_2_(*v*), *k*_3_(*v*)) is obtained from the algorithm in Section 5.3, and the tree is obtained *k*_1_(*v*), *k*_2_(*v*), *k*_3_(*v*)(of its three subtrees appear, followed by the trifurcating tree associated with rank *v*. by recursive application of the algorithm.

### 5.4 Special case: at-most-trifurcating trees

As an example of an at-most-*k*-furcating ranking, we consider *at-most-trifurcating* trees, in which internal nodes are permitted to bifurcate or trifurcate. *An* becomes the set of at-most-trifurcating unlabeled rooted trees with *n* leaves. For *n* = 1, *An* contains the unlabeled tree with one leaf. For *n* = 2, *An* contains the unique unlabeled tree with two leaves. We consider *k* = 3 in Definition 5.2.

#### Definition 5.4.

The ranking *f* : *A* → ℤ^+^ for at-most-trifurcating unlabeled rooted trees is a function that satisfies

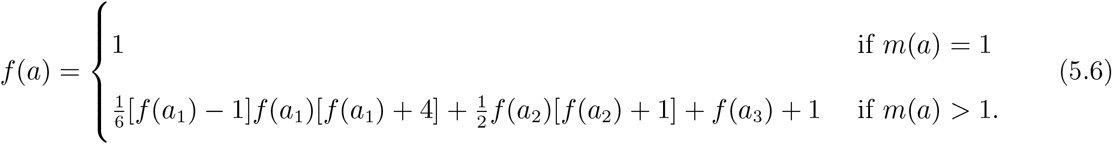

#### Theorem 5.5.

The function *f* ^−1^ : ℤ^+^ → *A* gives the three coordinates of the tree whose rank is *v*, and it satisfies

a. *f* ^−1^(1) is the tree with one leaf, and
b. for *v* ℤ 2, *f* ^−1^(*v*) is the tree *t* ∈ *T* whose subtrees have the ranks:

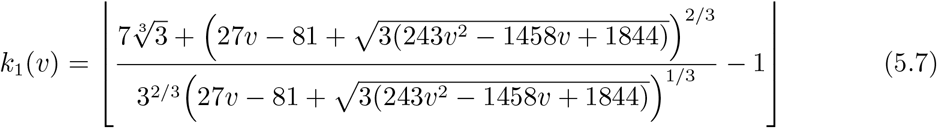

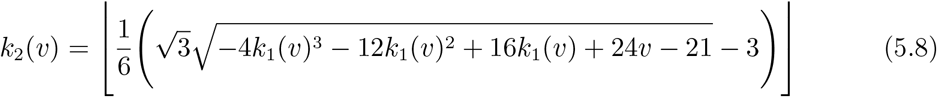

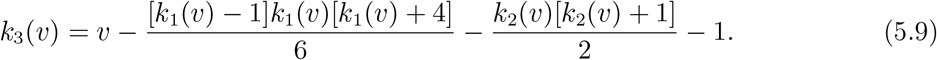

*Proof*. Given *v* ℤ 2, we seek to find the unique *f* (*a*_1_), ℤ *f* (*a*_2_), *f* (*a*_3_) with *f* (*a*_1_) ⩾ *f* (*a*_2_) *f* (*a*_3_), *f* (*a*_1_) ⩾ 1, *f* (*a*_2_) 1, and *f* (*a*_3_) ℤ 0, that solves

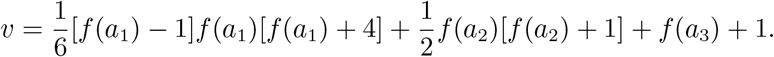

The solution for *f* (*a*_1_) in eq. 5.7 is obtained by solving the inequality

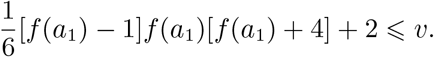

Next, the equation for *f* (*a*_2_) in eq. 5.8 is obtained by solving

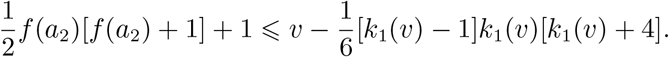

Finally, the inverse function for the third coordinate is given by eq. 5.9.

## 6 Discussion

We have devised ranking systems for multifurcating unlabeled trees, considering trees for which each internal node contains a fixed number of child nodes *k* as well as trees for which each internal node contains *at most k* child nodes. The general rankings extend a ranking for bifurcating trees, relying on lexicographic orders for subtrees, an analogy between subtree ranks and “place value,” and Macaulay’s binomial expansion. The main results are summarized in Table 1.

**Table 1:**
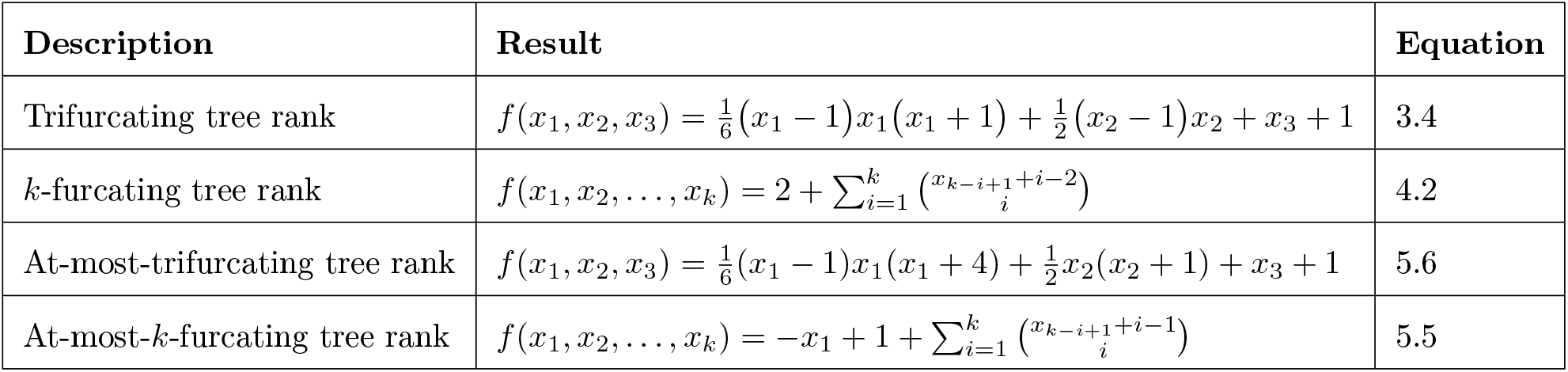
Summary of main results

| Description | Result | Equation |
| --- | --- | --- |
| Trifurcating tree rank | $f(x_1, x_2, x_3) = \frac{1}{6}(x_1 - 1)x_1(x_1 + 1) + \frac{1}{2}(x_2 - 1)x_2 + x_3 + 1$ | 3.4 |
| $k$-furcating tree rank | $f(x_1, x_2, \dots, x_k) = 2 + \sum_{i=1}^k \binom{x_{k-i+1}+i-2}{i}$ | 4.2 |
| At-most-trifurcating tree rank | $f(x_1, x_2, x_3) = \frac{1}{6}(x_1 - 1)x_1(x_1 + 4) + \frac{1}{2}x_2(x_2 + 1) + x_3 + 1$ | 5.6 |
| At-most-$k$-furcating tree rank | $f(x_1, x_2, \dots, x_k) = -x_1 + 1 + \sum_{i=1}^k \binom{x_{k-i+1}+i-1}{i}$ | 5.5 |

For the *k*-furcating trees with *k* ⩾ 3, as the rank *v* increases, in comparison with the bifurcating case (*k* = 2), we observe faster growth in the number of leaves of *k*-furcating trees associated with rank *v* (Figures 1-3). Because each internal node necessarily has *k* descendants, relatively few multifurcating trees possess small numbers of leaves. Thus, the number of leaves of *k*-furcating trees increases quickly with *v*. This effect is magnified as *k* increases, so that for large *k*, the number of leaves in the tree with rank *v* is generally larger than for small *k*.

For the at-most-*k*-furcating trees with *k* ⩾ 3, the growth in the number of leaves is not as fast as for *k*-furcating trees (Figures 1, 4, and 5). We can explain this observation by noting that for at-most-*k*-furcating trees, the flexibility in the number of child nodes that could descend from an internal node increases the number of possible trees for a given number of leaves, in comparison with the *k*-furcating case. As *k* increases for at-most-*k*-furcating trees, more possible trees with a given number of leaves exist in the lexicographical ordering, so that the trees in a lexicographical ordering associated with a lower *k* value are contained in the ordering for a higher *k* value. Hence, as *k* increases, the number of leaves in the tree with fixed rank *v* tends to decrease.

**Figure 5:**
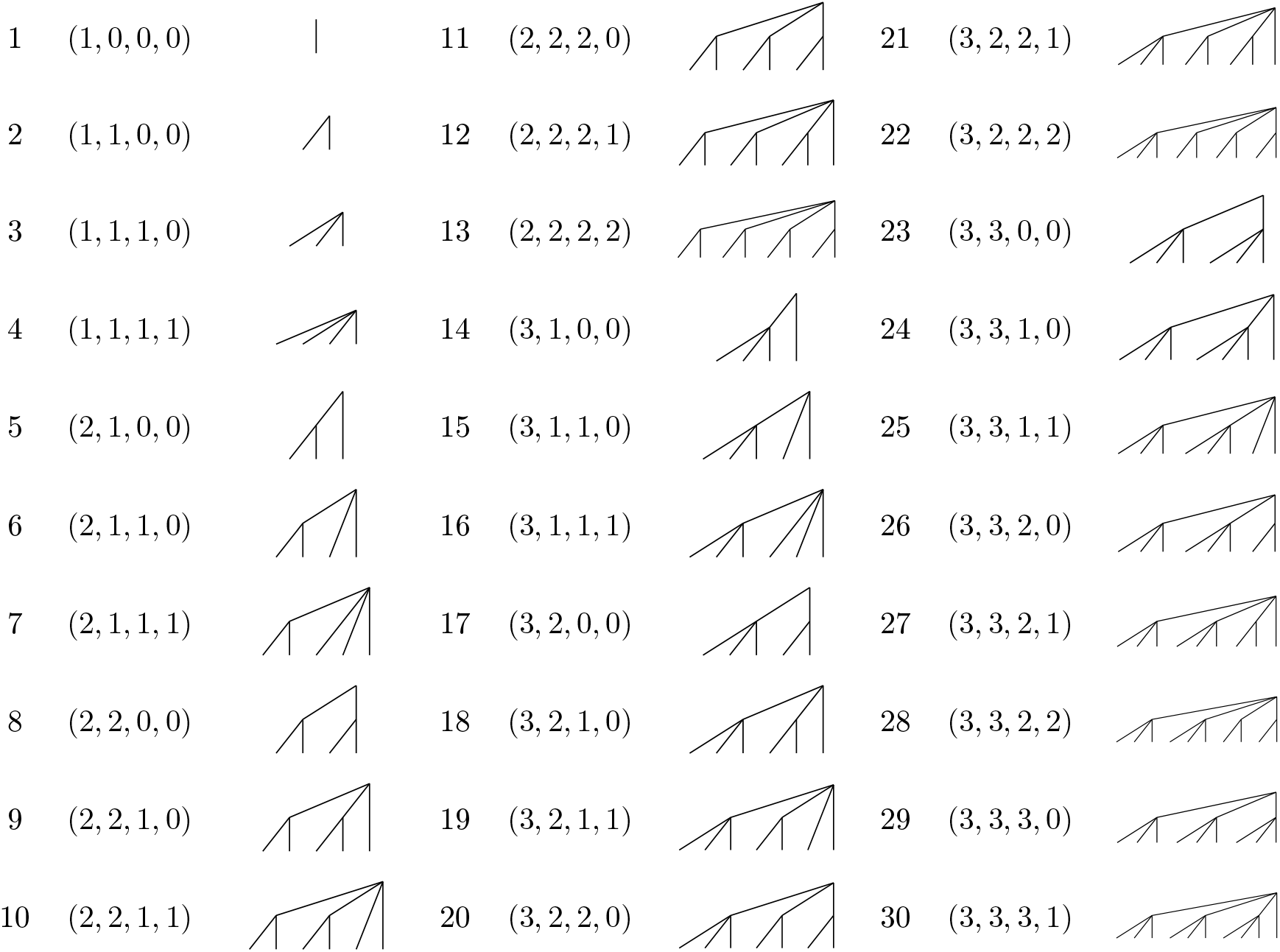
At-most-4-furcating trees associated with specified ranks. For each rank *v* from 1 to 30, the ranks The ordered quadruple (*k*_1_(*v*), *k*_2_(*v*), *k*_3_(*v*), *k*4(*v*)) is obtained from the algorithm in Section 5.3, and the tree (*k*_1_(*v*), *k*_2_(*v*), *k*_3_(*v*), *k*4(*v*)) of its four subtrees appear, followed by the 4-furcating tree associated with rank *v*. is obtained by recursive application of the algorithm.

The rankings for multifurcating unlabeled trees have potential uses in phylogenetic studies. In the analysis of phylogenetic trees, particularly in the context of the dynamics of pathogen sequences during epidemics, the unlabeled shape of a tree can provide insight about features of an evolutionary process [1], [5]. For rapidly spreading epidemics, a pathogen sequence in one infected individual can branch into multiple sequences in multiple subsequently infected individuals before mutations begin to accumulate in the subsequent sequences. Hence, phylogenetic trees during epidemics often appear to possess a multifurcating structure. The ranking schemes for multifurcating trees can potentially be used in statistics that are informative about the epidemic process; as is the case for bifurcating trees, they may be possible to use in “tree balance” concepts for multifurcating trees [3]. We note that our conceptualization of multifurcating trees differs from some that have been previously considered. Colijn & Plazzotta [1] commented on extending their enumeration scheme from bifurcating to multifurcating trees. However, their scheme for multifurcating trees permitted internal nodes with only a single descendant, a somewhat unnatural choice for mathematical phylogenetics. In enumerating multifurcating trees, Felsenstein [2, p. 33] fixed the number of leaves at *n* and counted multifurcating trees with at most *n* leaves. This calculation corresponds in our scheme to the enumeration of at-most-*n*-furcating trees with at most *n* leaves. However, in our ranking, many trees with *more than n* leaves would be assigned ranks smaller than the total number of at-most-*n*-furcating trees with at most *n* leaves—so that Felsenstein’s set of trees does not naturally correspond to a set of trees tabulated by any of our schemes. As mathematical phylogenetic analysis continues to examine multifurcating trees from the rapidly diversifying sequences that arise during the spread of pathogenic organisms, it will be of interest to clarify which definitions for sets of multifurcating trees give rise to the greatest potential for meaningful biological interpretation.

## Acknowledgements

We acknowledge NSF grant BCS-2116322 for support.

